# Read-Based Phasing of Related Individuals

**DOI:** 10.1101/037101

**Authors:** Shilpa Garg, Marcel Martin, Tobias Marschall

## Abstract

**Motivation:** Read-based phasing deduces the haplotypes of an individual from sequencing reads that cover multiple variants, while genetic phasing takes only genotypes as input and applies the rules of Mendelian inheritance to infer haplotypes within a pedigree of individuals. Combining both into an approach that uses these two independent sources of information - reads and pedigree - has the potential to deliver results better than each individually.

**Results:** We provide a theoretical framework combining read-based phasing with genetic haplotyping, and describe a fixed-parameter algorithm and its implementation for finding an optimal solution. We show that leveraging reads of related individuals jointly in this way yields more phased variants and at a higher accuracy than when phased separately, both in simulated and real data. Coverages as low as 2× for each member of a trio yield haplotypes that are as accurate as when analyzed separately at 15× coverage per individual.

**Availability:** https://bitbucket.org/whatshap/whatshap (branch pedmec)

## 1 Introduction

Humans are diploid and determining the sequences of both homologous copies of each chromosome is desirable for many reasons. These two sequences per chromosome are known as *haplotypes.* The process of obtaining them is known as *haplotyping* or *phasing*, two terms that we will use interchangeably. Haplotype-resolved genetic data can be used, for instance, for population genetic analyses of admixture, migration, and selection, but also to study allele-specific gene regulation, compound heterozygosity, and their roles in human disease. We refer the reader to Tewhey *et al.* (2011) and Glusman *et al.* (2014) for detailed reviews on the relevance of haplotyping.

### General Approaches for Haplotyping

There are three major approaches to phasing. First, haplotypes can be inferred from genotype information of large cohorts based on the idea that common ancestry gives rise to shared haplotype tracts, as reviewed by Browning and Browning (2011). This approach is known as *statistical* or *population-based phasing*. It can be applied to unrelated individuals and only requires genotype data, which can be measured at low cost. While very powerful for common variants, this technique is less accurate for phasing rare variants and cannot be applied at all to private or *de novo* variants. Second, haplotypes can be determined based on genotype data of related individuals, known as *genetic haplotyping* (Glusman *et al.*, 2014). To solve the phasing problem, one seeks to explain the observed genotypes under the constraints imposed by the Mendelian laws of inheritance, while being parsimonious in terms of recombination events. For larger pedigrees, such as parents with many children, this approach yields highly accurate phasings (Roach *et al.*, 2011). On the other hand, it is less accurate for single mother-father-child trios and has the intrinsic limitation of not being able to phase variants that are heterozygous in all individuals. Third, the sequences of the two haplotypes can be determined experimentally, called *molecular haplotyping*. Many techniques do not resolve the full-length haplotypes but yield blocks of varying sizes. Approaches furthermore largely differ in the amount of work, DNA, and money they require. On one end of the scale, next-generation sequencing (NGS) instruments generate local phase information of the length of a sequenced fragment at ever-decreasing costs. Another approach consists in breaking both homologous chromosomes into (larger) fragments and separating them into a number of pools such that each pool is unlikely to contain fragments from the same locus of both haplotypes. This can, for instance, be achieved by dilution followed by bar-coded short-read sequencing. To achieve molecular haplotyping over the range of a full chromosome, protocols have been invented to physically separate the two homologous chromosomes, for example by microscopy-based chromosome isolation, fluorescence-activated sorting, or microfluidics-based sorting. These and other experimental techniques for molecular haplotyping have been surveyed by Snyder *et al.* (2015). They are of great interest because they facilitate phasing of rare variants for single individuals. Rare variants have been postulated to contribute considerably to clinical traits and are hence of major interest.

### Haplotype Assembly

When many haplotype fragments are available for one individual, for instance from sequencing, one can attempt to reconstruct the full haplotypes or at least to obtain larger blocks. This process is known as *haplotype assembly, single-individual haplotyping*, or *read-based phasing* (in case the fragments indeed stem from sequencing reads). It requires reads that span two or more heterozygous variants. In order to be successful, reads covering as many pairs of consecutive heterozygous variants as possible are desirable. At present, third generation sequencing platforms, as marketed by Pacific Biosciences (PacBio) and Oxford Nanopore, become more widespread and offer reads spanning thousands to tens of thousands of nucleotides. Although error-rates are much higher than for common second generation technologies, the longer reads provide substantially more phase information and hence render them promising platforms for read-based phasing.

We adapt the common assumption that all variants to be phased are bi-allelic and non-overlapping. Then, the input to the haplotype assembly problem can be specified by a matrix with entries from {0,1, –}, hav-ing one row per read and one column per variant. Entries of 0 or 1 indicate that the read in that row gives evidence for the reference or alternative allele for the variant in that column, respectively. An entry of “–” signifies that a variant is not covered by that read. In case all reads are error-free and mapped to correct positions, the set of rows in that matrix admits a bipartition such that each of the two partitions is *conflict free*. Here, two rows are defined to be conflicting if they exhibit different non-dash values in the same column; that is, one entry is 0 and the other one is 1. If such a bipartition exists, the matrix is called *feasible*.

To formalize the haplotype assembly problem in the face of errors, we define *operations* on the input matrix and ask for the minimum number of operations one needs to apply to render it feasible. Different such operations have been studied, in particular removal of rows, resulting in the *Minimum Fragment Removal (MFR)* problem, removal of columns, resulting in the *Minimum SNP Removal (MSR)* problem, and flipping of bits, resulting in the *Minimum Error Correction (MEC)* problem. All three problems are NP-hard (Lancia *et al.*, 2001; Cilibrasi *et al.*, 2007). Flipping of bits corresponds to correcting sequencing errors and hence the MEC problem has received most attention in the literature and is most relevant in practice. A wealth of exact and heuristic approaches to solve the MEC problem exists. Exact approaches, which solve the problem optimally, include integer linear programming (Fouilhoux and Mahjoub, 2012; Chen *et al.*, 2013b), and fixed-parameter tractable (FPT) algorithms (He *et al.*, 2010; Patterson *et al.*, 2015; Pirola *et al.*, 2015). Refer to the reviews by Schwartz (2010) and Rhee *et al.* (2015) for further related approaches.

Here, we build upon our previous approach What-sHap (Patterson *et al.*, 2014, 2015) and generalize it to jointly handle sequencing reads of related individuals. WhatsHap is an FPT approach that solves the (weighted) MEC problem optimally in time exponential in the maximum coverage, but linear in the number of variants. In particular the runtime does not explicitly depend on the read length. These properties make it particularly apt for current long-read data. This has also been observed by Kuleshov (2014), who approached the weighted MEC problem in a message-passing framework and, by doing so, independently arrived at the same DP algorithm used in WhatsHap. The exponential runtime in the maximum coverage does not constitute a problem in practice because reads can be removed in regions of excess coverage without loosing much information. The evaluation by Patterson *et al.* (2015) suggests that pruning data to a maximum coverage of 15× yields excellent results while an even higher coverage does not deliver a big additional improvement.

### Hybrid Approaches

The ideas underlying population-based phasing, genetic haplotyping, and read-based phasing have been combined in many ways to create hybrid methods. Delaneau *et al.* (2013), for instance, use local phase information provided by sequencing reads to enhance their population-based phasing approach SHAPEIT. Exploiting pedigree information for statistical phasing has also been demonstrated to significantly improve the inferred haplotypes (Marchini *et al.*, 2006; Chen *et al.*, 2013a). Using their heuristic read-based phasing approach HapCompass, Aguiar and Istrail (2013) note that combining reads from parent-offspring duos increases performance in regions that are identical by descent (IBD). Beyond this approach, we are not aware of prior work to leverage family information towards read-based phasing.

### Contributions

Here, we introduce a unifying formal framework to fully integrate read-based and genetic haplotyping. To this end, we define the Weighted Minimum Error-Correction on Pedigrees Problem, termed PedMEC, which generalizes the (weighted) MEC problem and accounts for Mendelian inheritance and recombination. This problem is NP-hard. We generalize the WhatsHap algorithm for solving this problem optimally and thereby show that PedMEC is fixed-parameter tractable. When the maximum coverage is bounded, the runtime of our algorithm is linear in the number of variants and does not explicitly depend on the read length, hence inheriting the favorable properties of WhatsHap.

We target an application scenario where related individuals are sequenced using error-prone long-read technologies such as PacBio sequencing. As a driving question motivating this research, we ask how much coverage is needed for resolving haplotypes in related individuals as opposed to single or unrelated individuals. Our focus is on phasing and we do not consider the genotyping step, which can either be done from the same data or from orthogonal and potentially cheaper data sources such as microarrays or short-read sequencing. On simulated and real PacBio data, we show that sequencing each individual in a mother-father-child trio to 5× coverage is sufficient to establish a high-quality phasing. This is in stark contrast to state-of-the-art single-individual read-based phasing, which yields worse results even for 15× coverage with respect to both error rates and numbers of phased variants. We furthermore demonstrate that our technique also exhibits favorable properties of genetic haplotyping approaches: Because of genotype relationships between related individuals, we are able to infer correct phases even *between* haplotype blocks that are *not connected* by any sequencing reads in any of the individuals.

### Example

Figure 1 shows seven SNP positions covered by reads in three related individuals. It illustrates how the ideas of genetic and read-based haplotyping complement each other. All genotypes at SNP 3 are heterozygous. Thus, its phasing cannot be inferred by genetic phasing, that is, using only the given genotypes and not the reads. SNP 4, in contrast, is not covered by any read in the child. When only using reads in the child (corresponding to single-individual read-based phasing), no inference can be made about the phase of SNP 4 and neither about the phase between SNP 3 and SNP 5. By observing that all seven child genotypes are compatible with the combination of brown and green haplotypes from the parents, however, these phases can be easily inferred. This example demonstrates that jointly using pedigree information, genotypes, and sequencing reads is very powerful for establishing phase information.

**Figure 1:**
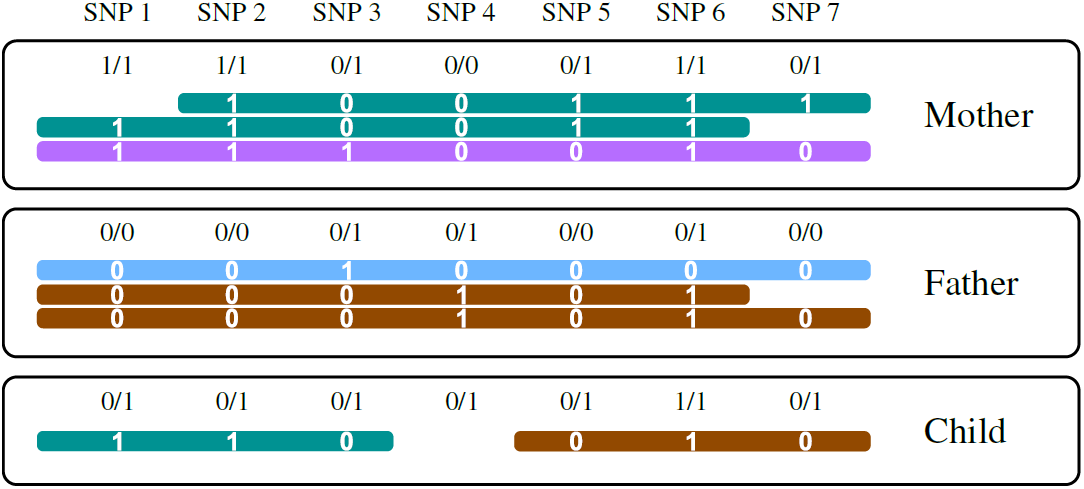
Seven SNP loci covered by reads (horizontal bars) in three individuals. Unphased genotypes are indicated by labels 0/0, 0/1, and 1/1. The alleles that a read supports are printed in white.

## 2 The Weighted Minimum Error Correction Problem on Pedigrees

Thus far, read-based phasing has predominantly been formulated as the Minimum Error Correction (MEC) problem (Cilibrasi *et al.*, 2007) and its weighted sibling wMEC (Greenberg *et al.*, 2004). We first re-state these problems formally to introduce our notation and then proceed to generalizing them to pedigrees.

The input to the MEC problem consists of a SNP matrix ℱ ∈ {0,1,–}^R×M^, where R is the number of reads and M is the number of variants along a chromosome. Each matrix entry ℱ (*j, k*) is 0 (indicating that the read matches the reference allele) or 1 (indicating that the read matches the alternative allele) if the read covers that position and “–” otherwise. Note that the “–” character can also be used to encode the unsequenced “internal segment” of a paired-end read.

### Definition 1 (Distance)

*Given two vectors r*_1_, *r*_2_ ∈ {0, 1, –}^*M*^, *the* distance d(*r*_1_, *r*_2_) *between r*_1_ *and r*_2_ *is given by the number of mismatching non-dash characters. Formally*,

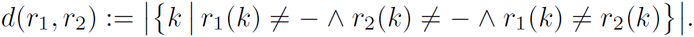

### Definition 2 (Feasibility)

*A SNP matrix* ℱ ∈ {0, 1, –}^*R×M*^ *is called* feasible *if there exists a bipartition of rows (i. e., reads) into two sets such that all pairwise distances of two rows within the same set are zero.*

Feasibility of a matrix ℱ is equivalent with the existence of two haplotypes *h*^0^, *h*^1^ ∈ {0,1, –}^*M*^ such that every read *r* in the matrix has a distance of zero to *h*^0^ or to *h*^1^ (or both). The MEC problem can now simply be stated in terms of flipping bits in ℱ, where entries that are 0 or 1 can be flipped and “–” entries are fixed.

### Problem 1 (MEC)

*Given a matrix* ℱ ∈ {0, 1, –}^*R*×*M*^, *flip a minimum number of entries in* ℱ *to obtain a feasible matrix*.

The MEC problem is NP-hard (Cilibrasi *et al.*, 2007). The weighted version of the problem associates a cost to every matrix entry. This is useful since each nucleotide in a sequencing read usually comes with a “phred-scaled” base quality *Q* that corresponds to an estimated probability of 10^-*Q*/10^ that this base has been wrongly sequenced. These phred scores can hence serve as costs of flipping a letter, allowing less confident base calls to be corrected at lower cost compared to high confidence ones.

### Problem 2 (wMEC)

*Given a matrix* ℱ ∈ {0, 1, –}^*R*×*M*^ *and a weight matrix* 𝒲 ∈ ℕ^*R*×*M*^, *flip entries in ℱ to obtain a feasible matrix, while minimizing the sum of incurred costs, where flipping entry* ℱ (*j*, *k*) *incurs a cost of* 𝒲 (*j*, *k*).

We now generalize wMEC to account for multiple individuals in a pedigree simultaneously while modelling inheritance and recombination. We assume our pedigree to contain a set of *N* individuals *𝓘* = {1,…, *N*}. Relationships between individuals are given as a set of (ordered) mother-father-child triples 𝒯. For example, if *𝓘* = {1, 2, 3, 4}, then 𝒯 = {(1, 2, 3), (1, 2,4)} corresponds to a pedigree where individuals 1 and 2 are the parents of individuals 3 and 4. We only consider non-degenerate cases without circular relationships and where each individual appears as a child in at most one triple. Furthermore, we assume all considered variants to be non-overlapping and bi-allelic. Each individual *i* comes with a *genotype vector g_i_* ∈ {0,1, 2}^*M*^, giving the genotypes of all *M* variants. Genotypes 0, 1, and 2 correspond to being homozygous in the reference allele, heterozygous, and homozygous in the alternative allele, respectively. In the context of phasing, we can restrict ourselves to the set of variants that are heterozygous in at least one of the individuals, that is, to variants *k* such that *g_i_* (*k*) = 1 for at least one individual *i* ∈ *𝓘*. For each individual *i* ∈ *𝓘*, a number of *R_i_* aligned sequencing reads is provided as input, giving rise to one SNP matrix ℱ*_i_* ∈ {0,1, –}^*R*×*M*^ and one weight matrix 𝒲_*i*_ ∈ ℕ^*R_i_*×*M*^ per individual. We seek to compute two haplotypes 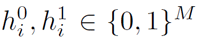 for all individuals *i* ∈ *𝓘*. As before in the MEC problem, we want these haplotypes to be consistent with the sequencing reads.

In addition, we want the haplotypes to respect the constraints given by the pedigree. Recall that in each parent, the two homologous chromosomes recombine during meiosis to give rise to a haploid gamete that is passed on to the offspring. Therefore, each haplotype of a child should be representable as a mosaic of the two haplotypes of the respective parent with few recombination events. To control the number of recombination events, we assume a per-site recombination cost of 𝒳(*k*) to be provided as input. Controlling the recombination cost per site is important because it is not equally likely to happen at all points along a chromosome. Instead *recombination hotspots* exist, where recombination is much more likely to occur (and should hence be penalized less strongly in our model). The cost 𝒳(*k*) should be interpreted as the (phred-scaled) probability that a recombination event occurs between variant *k*–1 and variant *k*. To formalize the inheritance process, we define *transmission vectors* 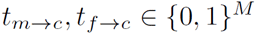 for each triple (*m*, *f*, *c*) ∈ *T*. The values *t_m⟶c_*(*k*) and *t_f⟶c_*(*k*) tell which allele at site *k* is transmitted by mother and father, respectively. The haplotypes we seek to compute have to be *compatible* with transmission vectors, defined formally as follows.

### Definition 3 (Transmission vector compatibility)

*For a given trio* (*m*, *f*, *c*) ∈ 𝒯, *the haplotypes* 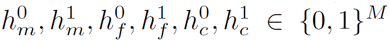 *are* compatible *with the transmission vectors* 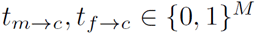 *if*

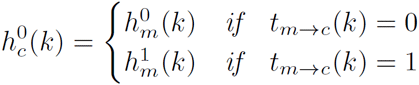

*and*

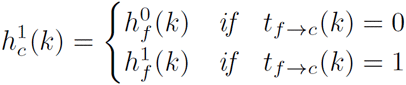

*for all k* ∈ {1,…, *M*}.

With this notion of transmission vectors, recombination events are characterized by changes in the transmission vector, that is, by positions *k* with *t_m⟶c_*(*k*–1) ≠ *t_m⟶c_*(*k*) or *t_f⟶c_*(*k*–1) ≠ *t_f⟶c_*(*k*). Given our recombination cost vector 𝒳, the cost associated to a transmission vector can be written as follows (in slight abuse of notation).

### Definition 4 (Transmission cost)

*For a transmission vector t_p⟶c_* ∈ {0, 1}^*M*^ *with p* ∈ {*m*, *f*} *and a recombination cost vector* 𝒳 ∈ ℕ^*M*^, *the cost of t_p⟶c_ is defined as*

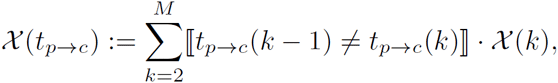

*where* ⟦S⟧ = 1 *if statement S is true and* 0 *otherwise*.

To state the problem of jointly phasing all individuals in *𝓘* formally, it is instrumental to consider the set of matrix entries to be flipped explicitly. We will therefore introduce a set of index pairs E_*i*_ ⊂ {1,…, *R_i_*} × {1,…, *M*} where (*j*, *k*) ∈ E_*i*_ if and only if the bit in row *j* and column *k* of matrix ℱ_*i*_ is to be flipped.

### Problem 3 (Weighted Minimum Error Correction on Pedigrees, PedMEC)

*Let a set of individuals 𝓘* = {1, …, *N*}, *a set of relationships* 𝒯 *on 𝓘*, *recombination costs* 𝒳 ∈ ℕ^*M*^, *and*, *for each individual i* ∈ *I*, *a sequencing read matrix* ℱ_*i*_ ∈ {0, 1, –}^*R_i_*^×^*M*^ *and corresponding weights* 𝒲_*i*_ ∈ ℕ^*R_i_*×*M*^ *be given. Determine a set of matrix entries to be flipped E_i_* ⊂ {1, …, *R_i_*} × {1,…, *M*} *to make* ℱ_*i*_ *feasible and two haplotypes* 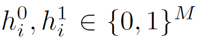 *for each individual i* ∈ *𝓘 as well as two transmission vectors t_m⟶c_, t_f⟶c_* ∈ {0, 1}^*M*^ *for each trio* (*m, f, c*) ∈ 𝒯 *such that*

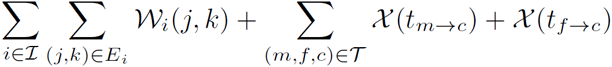

*takes a minimum, subject to the constraints that all haplotypes are compatible with the corresponding transmission vectors, if existing.*

Note that for the special case of *𝓘* = {1} and 𝒯 = ∅, PedMEC is identical to wMEC. Therefore, the PedMEC problem is also NP-hard. As discussed in Section 1, we are specifically interested in an application scenario were the genotypes are already known. By using genotype data, we aim to most beneficially combine the merits of genetic haplotyping and read-based haplotyping. We therefore extend the PedMEC problem to incorporate genotypes and term the resulting problem PedMEC-G.

### Problem 4 (PedMEC with genotypes, PedMEC-G)

*Let the same input be given as for Problem 3 (PedMEC) and, additionally, a genotype vector g_i_* ∈ {0, 1, 2}^*M*^ *for each individual i* ∈ *𝓘*. *Solve the Ped-MEC problem under the additionalconstraints that* 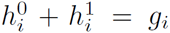 *for all i* ∈ *𝓘*, *where* “+” *refers to acomponent-wise addition ofvectors*.

For the classical MEC problem, additionally assuming that all sites to be phased are heterozygous is common (Chen *et al.*, 2013b). This variant of the MEC problem is a special case of PedMEC-G with *𝓘* = {1} and 𝒯 = ∅ and *g*_1_ (*k*) = 1 for all *k*.

## 3 Algorithm

### Solving MEC and wMEC

WhatsHap (Patterson *et al.*, 2015) is a dynamic programming (DP) algorithm to optimally solve the wMEC problem. It runs in 𝒪(2^c^. *M*) time, where *M* is the number of variants to be phased and c is the maximum physical coverage (which includes internal segments of paired-end reads). The general idea is to proceed column-wise from left to right while maintaining a set of active reads. Each read remains active from its first non-dash position to its last non-dash position in ℱ. Let the set of active reads in column *k* be denoted by *A(k)*. Note that *c* = max_*k*_{*A(k)*}. For each column *k* of ℱ, we fill a DP table column *C*(*k*,.) with 2^|*A(k)*|^ entries, one entry for each bipartition *B* of the set of active reads *A(k)*. Each entry *C(k, B)* is equal to the cost of solving wMEC on the partial matrix consisting of columns 1 to *k* of ℱ under the assumption that the sought bipartition of the full read set *A*(1) ∪ … ∪ *A*(*k*) *extends B* according to the below definition.

#### Definition 5 (Bipartition extension)

*For a given set A and a subset A' ⊂ A, a bipartition B* = (*P, Q*) *of A is said to* extend *a bipartition B'* = (*P', Q'*) *of A' if P' ⊂ P and Q' ⊂ Q*.

By this semantics of DP table entries *C(k, B)*, the minimum of the last column min_*B*_ {*C*(*M, B*)} is the optimal wMEC cost.

### Algorithm Overview: Solving PedMEC and PedMEC-G

In the following, we will see how this idea can be extended for solving PedMEC and PedMEC-G. The basic idea is to use the same technique on the union of the sets of active reads across all individuals *i ∈ 𝓘*, while adding some extra book-keeping to satisfy the additional constraints imposed by pedigree and genotypes. Let *A_i_(k)* be the set of active reads in column *k* of ℱ_*i*_. We now define *A(k)* = ⋃_*i∈𝓘*_*A_i_(k)*. A bipartition *B* = (*P,Q*) of *A(k)* now induces bipartitions for each individual: *B_i_* = (*P*∩*A_i_(k)*), *Q*∩*A_i_(k)*).

As before, we consider all bipartitions of *A(k)* for each column *k*, but now additionally distinguish between all possible transmission values. We assume the set of trio relationships 𝒯 to be (arbitrarily) ordered and use a tuple *t* ∈ {0,1}^2|𝒯|^ to specify an assignment of transmission values. Such an assignment *t* can later (during backtracing) be translated into the sought transmission vectors: Assuming *t* to be an optimal such tuple at column *k*, its relation to the transmission vectors is given by

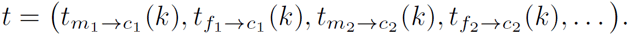

The transmission tuples give rise to one additional dimension of our DP table for PedMEC(-G), as compared to the DP table for wMEC. For each column *k*, we compute table entries *C(k, B,t)* for all 2^|*A(k)*|^ bipartitions of reads and all 2^2|𝒯|^ possible transmission tuples, for a total of 2^|*A(k)*|+2|𝒯|^ entries in this column.

### Computing Local Costs

Along the lines of Patterson *et al.* (2015), we first describe how to compute the cost incurred by flipping matrix entries in each column, denoted by Δ_*C*_(*k, B,t*), and then explain how to combine them with entries in *C*(*k* – 1, .,.) to compute the cost *C(k, B, t)*. The crucial point for dealing with reads from multiple individuals in a pedigree is to realize that matrix entries from haplotypes that are identical by descent (IBD) need to be identical (or need to be flipped to achieve this). For unrelated individuals (i.e. 𝒯 = ∅), none of the haplotypes are IBD, giving rise to 2|*𝓘*| sets of reads for the 2|*𝓘*| unrelated haplotypes. These 2|*𝓘*| sets of reads are given by B and the cost Δ_*C*_(*k, B,t)* can be computed by flipping all matrix entries of reads within the same set to the same value.

For a non-empty 𝒯, the transmission tuple *t* tells which parent haplotypes are passed on to which child. In other words, *t* identifies each child haplotype to be IBD to a specific parent haplotype. We can therefore merge the corresponding sets of reads since all reads coming from haplotypes that are IBD need to show the same allele and need to be flipped accordingly. In total, we obtain 2|*𝓘*|-2|𝒯 | sets of reads, since each trio relationship implies merging two pairs of sets. We write *S(k, B, t)* to denote this set of sets of reads induced by bipartition *B* and transmission tuple *t* in column *k*. The cost 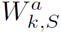 of flipping all entries in a read set *S ∈ S(k, B, t)* to the same allele *a* ∈ {0, 1} is given by

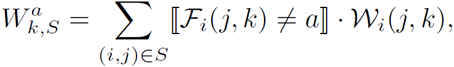

where we identify reads in *S* by a tuple (*i, j*), telling that it came from individual *i* and corresponds to row *j* in ℱ_*i*_. For PedMEC, i.e. if no constraints on genotypes are present, every set *S* can potentially be flipped to any allele *a* ∈ {0, 1}. Hence, the cost is given by

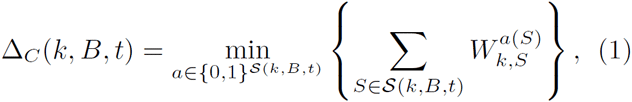

that is, we minimize the sum of costs incurred by each set of reads *S* ∈ *S(k, B, t)* over all possible assignments of alleles to read sets. For PedMEC-G, this minimization is constrained to only consider allele assignments consistent with the given genotypes. To ensure that valid assignments exist, we assume the input genotypes to be free of Mendelian conflicts.

### Computing a Column of Local Costs

To compute the whole column Δ_*C*_(*k*, .,.), we proceed as follows. In an outer loop, we enumerate all 2^2|𝒯|^ values of the transmission tuple *t*. For each value of *t*, we perform the following steps: We start with bipartition *B* = (*A(k)*, ∅) and compute all 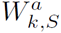 for all sets *S* ∈ *S (k, B, t)* and all *a* ∈ {0,1}, which can be done in 𝒪(|*A(k)*| + |*𝓘*|) time. Next we enumerate all bipartitions in Gray code order, as done previously (Patterson *et al.*, 2015). This ensures that only one read is moved from one set to another in each step, facilitating constant time updates of the values 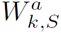. The value of Δ_*C*_(*k, B,t*) is then computed from the 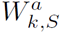’s according to Equation (1), which takes 𝒪(2^2|*𝓘*|^.|*𝓘*|) time. Computing the whole column Δ_*C*_(*k*, .,.) hence takes 𝒪(2^2^|𝒯|.(2^2|*A(k)*|^ + 2^2|*𝓘*|^.|*𝓘*|)) time.

### DP Initialization

The first column of the DP table, *C*(1, .,.), is initialized by setting *C*(1,*B,t*) := Δ_*C*_(1,*B,t*) for all bipartitions *B* and all transmission tuples *t*.

### DP Recurrence

Recall that *C(k, B, t)* is the cost of an optimal solution for input matrices restricted to the first *k* columns under the constraints that the sought bipartition extends *B* and that transmission happened according to *t* at site *k*. Entries in column *C*(*k* +1, .,.) should hence add up local costs incurred in column *k*+1 and costs from the previous column. To adhere to the semantics of *C*(*k*+1, *B, t*), only entries in column *k* whose bipartitions are *compatible* with *B* are to be considered as possible “predecessors” of *C*(*k* + 1, *B,t*).

#### Definition 6 (Bipartition compatibility)

*Let B = (P, Q) be a bipartition of A and B' = (P', Q') be a bipartition of A'. We say that B and B' are compatible, written B ≃ B', if P ⋂ (A ⋂ A') = P' ⋂ (A ⋂ A') and Q ⋂ (A ⋂ A') = Q' ⋂ (A ⋂ A')*.

Two bipartitions are therefore compatible when they agree on the intersection of the underlying sets. Besides ensuring that bipartitions are compatible, we need to incur recombination costs in case the transmission tuple *t* changes from *k* to *k* + 1. Formally, entries in column *k* + 1 are given by

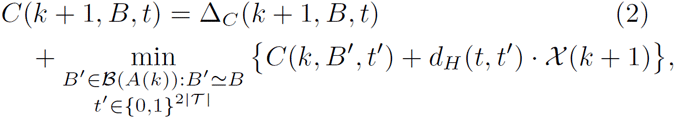

where ß (*A(k)*) denotes the set of all bipartitions of *A(k)* and *d_H_* is the Hamming distance. The distance *d_H_(*t, t'*)* hence gives the number of changes in transmission vectors and thus the term *d_H_(t,t')*.𝒳(*k* + 1) gives the recombination cost to be added.

### Projection Columns

To ease computing *C*(*k* + 1, *B, t*) via Equation (2), we use the same technique described by Patterson *et al.* (2015) and define intermediate *projection columns C*^⋂^(*k*, .,.). They can be thought of as being *between* columns *k* and *k* + 1. Consequently, they are concerned with bipartitions of the intersection of read sets *A(k)* ⋂ *A*(*k* + 1) and hence contain 2^|*A*(*k*)⋂*A*(*k*+1)| + 2|T|^ entries, which are given by

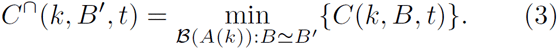

These projection columns can be created while computing *C*(*k*, .,.) at no extra (asymptotic) runtime. Using these projection columns, Equation (2) becomes

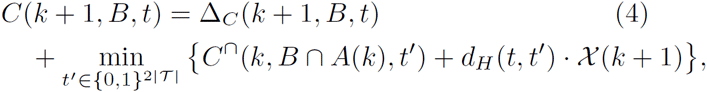

where *B ⋂ A(k)* := (*P ⋂ A(k)*, *Q ⋂ A(k)*) for *B* = (*P,Q*). We have therefore reduced the runtime of computing this minimum to 𝒪(2^2|𝒯|^).

### Runtime

Computing one column of local costs, Δ_*C*_(*k*, ...), takes 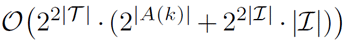 time, as discussed above. For each entry, we use Equation (4) to compute the aggregate value of cost incurred in present and past columns. Over all columns, we achieve a runtime of 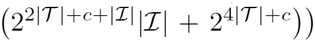, where *c* = max_*k*_ {|*A(k)*|} is the maximum coverage.

### Backtracing

An optimal bipartition and transmission vectors can be obtained by recording the indexes of the table entries that gave rise to the minima in equations (4) and (3) when filling the DP table and then backtracing starting from the optimal value in the last column. Optimal haplotypes are subsequently obtained using the bipartition and transmission vectors.

## 4 Experimental Setup

To evaluate the performance of our approach, we considered both real and simulated datasets.

### 4.1 Real Data

The Genome in a Bottle Consortium (GIAB) has characterized seven individuals extensively using eleven different technologies (Zook *et al.*, 2015). The data is publicly available. Here we consider the Ashkenazim trio, consisting of three related individuals: NA24143 (mother), NA24149 (father) and NA24385 (son). We obtained a consensus genotype call set (NIST_CallsIn2Technologies_05182015) provided by GIAB, containing variants called by two independent technologies. For our benchmark, we consider all bi-allelic SNPs on Chromosome 1 called in all three individuals, amounting to 141,256 in total, and use the provided (unphased) genotypes.

#### Ground Truth via Statistical Phasing

To generate a ground truth phasing for comparison, we used the population-based phasing tool SHAPEITv2-r837 (Delaneau *et al.*, 2014) with default parameters. The program was given the 1000 Genomes reference panel^1^, the corresponding genetic map^2^, and the unphased genotypes as input. SNPs present in the GIAB call set but absent in the reference panel were discarded, resulting in 140,744 phased SNPs for each individual, of which 58,551 were heterozygous in mother, 57,152 in father and 48,023 in child. We refer to this set of phased SNPs as *ground truth phased variants.* We emphasize that this phasing is solely based on genotypes and does not use phase information present in the reads in any way and hence is completely independent. Removing phase information from this call set results in *ground truth unphased genotypes*, which we use as input for read-based phasing experiments described below.

#### PacBio Data

For each individual, we downloaded aligned Pacific Biosciences (PacBio) reads^3^, which had an average coverage of 42.3× in mother, 46.8× in father and 60.2× in child, respectively. The average mapped read length across mother was 8,328 bp, father was 8,471 bp and child was 8,687 bp. For each individual, we separately downsampled the aligned reads to obtain data sets of 2×, 3×, 4×, 5×, 10×, and 15× average coverage.

### 4.2 Simulated Data

Despite the high-quality data set provided by GIAB, we sought to complement our experiments by a simulated data set. While the population-based phasing we use as ground truth is arguably accurate due to a large reference panel and the high-quality genotype data used as input, it is not perfect. Especially variants with low allele frequency present challenges for population-based phasers.

#### Virtual Child

As basis for our simulation, we use the haplotypes of the two parents from our ground truth phased dataset. We generated two haplotypes of a virtual child by applying recombination and Mendelian inheritance to the four parent haplotypes. In reality, recombination events are rare: All of Chromosome 1 spans a genetic distance of approximately 292 cM, corresponding to 2.9 expected recombination events along the whole chromosome. To include more recombinations in our simulated data set, we used the same genetic map as above, but multiplied recombination rates by 10. The recombination sites are sampled according to the probabilities resulting from applying Haldane’s mapping function to the genetic distances between two variants. In line with our expectation, we obtained 26 and 29 recombination sites for mother and father, respectively. The resulting child had 41,676 heterozygous variants.

#### Simulating PacBio Reads

We aimed to simulate reads that mimic the characteristics of the real PacBio data set as closely as possible. For this simulation, we incorporate the variants of each individual into the reference genome (hg19) to generate two true haplotypes for each individual in our triple. We used the PacBio-specific read simulator pbsim by Ono *et al.* (2013) to generate a 30× data set for Chromosome 1. The original GIAB reads were provided to pgsim as a template (via option `--sample-fastq`) to generate artificial reads with the same length profile. Next, we aligned the reads to the reference genome using BWA-MEM 0.7.12-r1039 by Li (2013) with option `-x pacbio`. As before for the real data, the aligned reads for each individual were downsampled separately to obtain data sets of 2×, 3×, 4×, 5×, 10×, and 15× average coverage.

### 4.3 Compared Methods

Our main goal is to analyze the merits of the PedMEC-G model in comparison to wMEC; in particular with respect to the coverage needed to generate a high-quality phasing. The algorithms to solve wMEC and PedMEC-G described in Section 3 have been implemented in the WhatsHap software package, distributed as Open Source software under the terms of the MIT license. The implementation of PedMEC-G is currently restricted to trios. It can be found in branch `pedmec` of the WhatsHap Git repository^4^. We emphasize that WhatsHap solves wMEC and PedMEC-G optimally. Since the focus of this paper is on comparing these two models, we do not include other methods for single-individual haplotyping. We are not aware of other trio-aware read-based phasing approaches that PedMEC-G could be compared to additionally.

The runtime depends exponentially on the maximum coverage. Therefore we prune the input data sets to a *target maximum coverage* using the read-selection method included in WhatsHap. This target coverage constitutes the only parameter of our method. For PedMEC-G, we prune the maximum coverage to 5× for each individual separately. For wMEC, we report results for 5× and 15× target coverage. The respective experiments are referred to as wMEC-5, PedMEC-G-5, and PedMEC-G-15. For wMEC, we use the additional “all heterozygous” assumption (see Section 3), to also give it the advantage of being able to “trust” the genotypes, as is the case for PedMEC-G. Both wMEC and PedMEC-G were provided with the ground truth unphased genotypes for the respective data set. PedMEC-G was additionally provided with the respective genetic map (original 1000G genetic map for real data and scaled by factor 10 for simulated data).

### 4.4 Performance Metrics

We compare each phased individual to the respective ground truth haplotypes separately and only consider sites heterozygous in this individual.

#### Phased SNPs

For read-based phasing of a single individual (wMEC), we say that two heterozygous SNPs are directly connected if there exists a read covering both. We compute the connected components in the graph where SNPs are nodes and edges are drawn between directly connected SNPs. Each connected component is called a *block*. For read-based phasing of a trio (PedMEC), we draw an edge when two SNPs are connected by a read in any of the three individuals. In both cases, we count a SNP as being *phased* when it is not the left-most SNP in its block (for the left-most SNP, no phase information with respect to its predecessors exists). All other SNPs are counted as *unphased*. Below, we report the average *fraction of unphased SNPs* over all three family members.

#### Phasing Error Rate

For each block, the first predicted haplotype is expressed as a mosaic of the two true haplotypes, minimizing the number of switches. This minimum is known as the *number of switch errors*. Note that the second predicted haplotype is exactly the complement of the first one, due to only considering heterozygous sites. When two switch errors are adjacent, they are subtracted from the number of switch errors and counted as one *flip error*. The *phasing error rate* is defined as the sum of switch and flip errors divided by the number of phased SNPs.

## 5 Results

We report the results of wMEC-5, wMEC-15, and PedMEC-G-5 for both data sets, *real* and *simulated*. All combinations of the three methods, two data sets, and six different average coverages (2×, 3×, 4×, 5×, 10× and 15×) were run. The predicted phasings are compared to the ground truth phasing for the respective data set. That is, for the real data set, we compare to the population-based phasing produced by SHAPEIT; for the simulated data set, we compare to the true haplotypes that gave rise to the simulated reads. Figure 2 shows the fraction of unphased SNPs in comparison to the phasing error rate (see Section 4.4) for all conducted experiments. A perfect phasing would be located in the bottom left corner.

**Figure 2:**
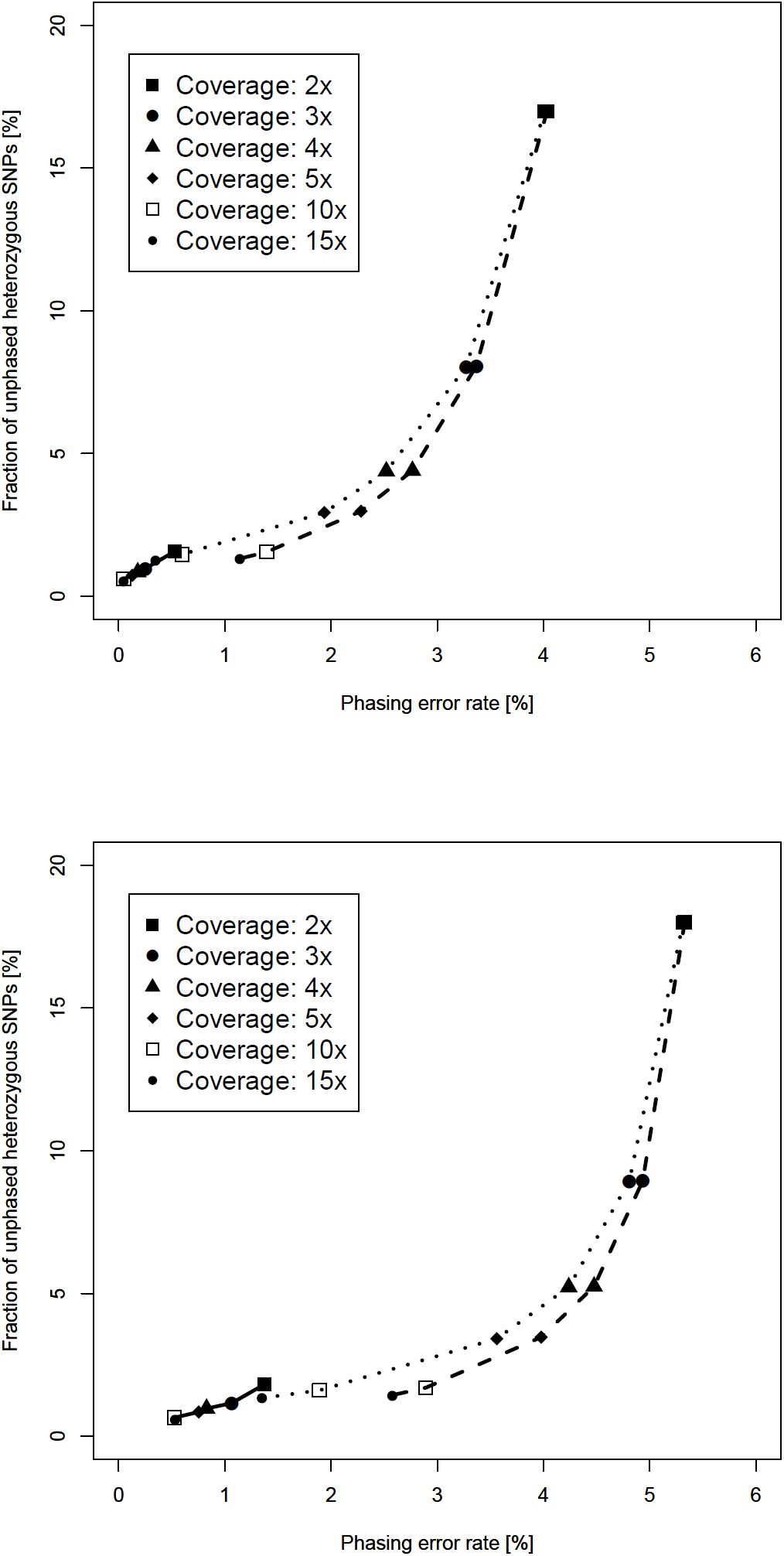
Simulated data set (top) and real dataset (bottom): phasing error rate (*x*-axis) versus completeness in terms of the fraction of unphased SNPs (*y*-axis) for PedMEC-G-5 (solid line), wMEC-5 (dashed line), and wMEC-15 (dotted line). Average coverage (per individual) of input data is encoded by different symbols.

### The Influence of Coverage

Increasing the average coverage is beneficial for phasing. For all three methods (wMEC-5, wMEC-15, and PedMEC-G-5) and both data sets, the phasing error rate and the fraction of unphased SNPs decrease monotonically when the average coverage is increased, as is clearly visible in Figure 2. The effect is much more drastic for wMEC than for PedMEC, however. Apparently, wMEC needs more coverage to compensate for PacBio’s high error rate while PedMEC can resort to exploiting family information to resolve uncertainty.

### The Value of Family Information

When operat-ing on the same input coverage, PedMEC-G-5 clearly outperformed wMEC-5 and even wMEC-15 in all cases tested. This was true for phasing error rates as well as for the fraction of phased positions. On the real data set with average coverage 10×, for instance, wMEC-5 and wMEC-15 reached an error rate of 2.9% and 1.9%, while it was 0.5% for PedMEC-G-5.

Most remarkingly, PedMEC-G-5 delivers excellent results already for very low coverages. When working with an average coverage as low as 2× for each family member, it achieves an error rate of 1.4% and a fraction of unphased SNPs of 1.8% (on real data). In contrast, wMEC-15 needs 15× average coverage on each individual to reach similar values (1.4% error rate and 1.3% unphased SNPs). When running on 5× data, PedMEC-G-5’s error rate and fraction of unphased positions decrease to 0.75% and 0.85%, respectively. Therefore, it reaches better results while requiring only a third of the sequencing data, which translates into significantly reduced sequencing costs.

### Comparison of Real and Simulated Data

When comparing results for simulated and real data, i.e. top and bottom plots in Figure 2, the curves appear similar, with some important difference. In terms the fraction of phased SNPs (*y*-axis), results are virtually identical. This indicates that our simulation pipeline establishes realistic conditions regarding this aspect. Differences in terms of error rates (*x*-axis) are larger. In general, error rates in the real data are larger than in the simulated data, which might be partly caused by a too optimistic error model during read simulation. On the other hand, the population-based phasing used as ground truth for the real data set will most likely also contain errors. Especially low-frequency variants present difficulties for population-based phasers. Given the fact that PedMEC-G-5’s performance does not seem to strongly depend on the input coverage and performs extremely well on simulated data, one might suspect that a significant fraction of the differences are indeed caused by errors in the ground truth phasing. For PedMEC-G-5 on 15× data, for example, we observe an error rate of 0.04% on simulated and 0.53% on real data.

### Phase Information Beyond Block Boundaries

Genetic phasing operates on genotypes of a pedigree, without using any sequencing reads. Figure 3 illustrates a case where we have two blocks that are not connected by reads in any individual. Nonetheless phase information can be inferred from the genotypes: Each block contains a SNP that is homozygous in both parents and heterozygous in the child, which immediately establishes which haplotype is maternal and which is paternal in both blocks. Note that this, in turn, also implies the phasing of the parents. By design, PedMEC-G implicitly exploits such information. To demonstrate this, we used the real data set and merged all blocks reported by PedMEC-G into one chromosome-wide block and determined the fraction of cases where phases were correctly inferred between blocks—and hence between two SNPs that are not connected by reads in any individual. This resulted in a fraction of 89.7% correctly inferred phased (averaged over all individuals and coverages; standard deviation 1.4%). Repeating the same for wMEC yielded 50.4% correctly inferred phased, as expected equalling a coin flip.

**Figure 3:**
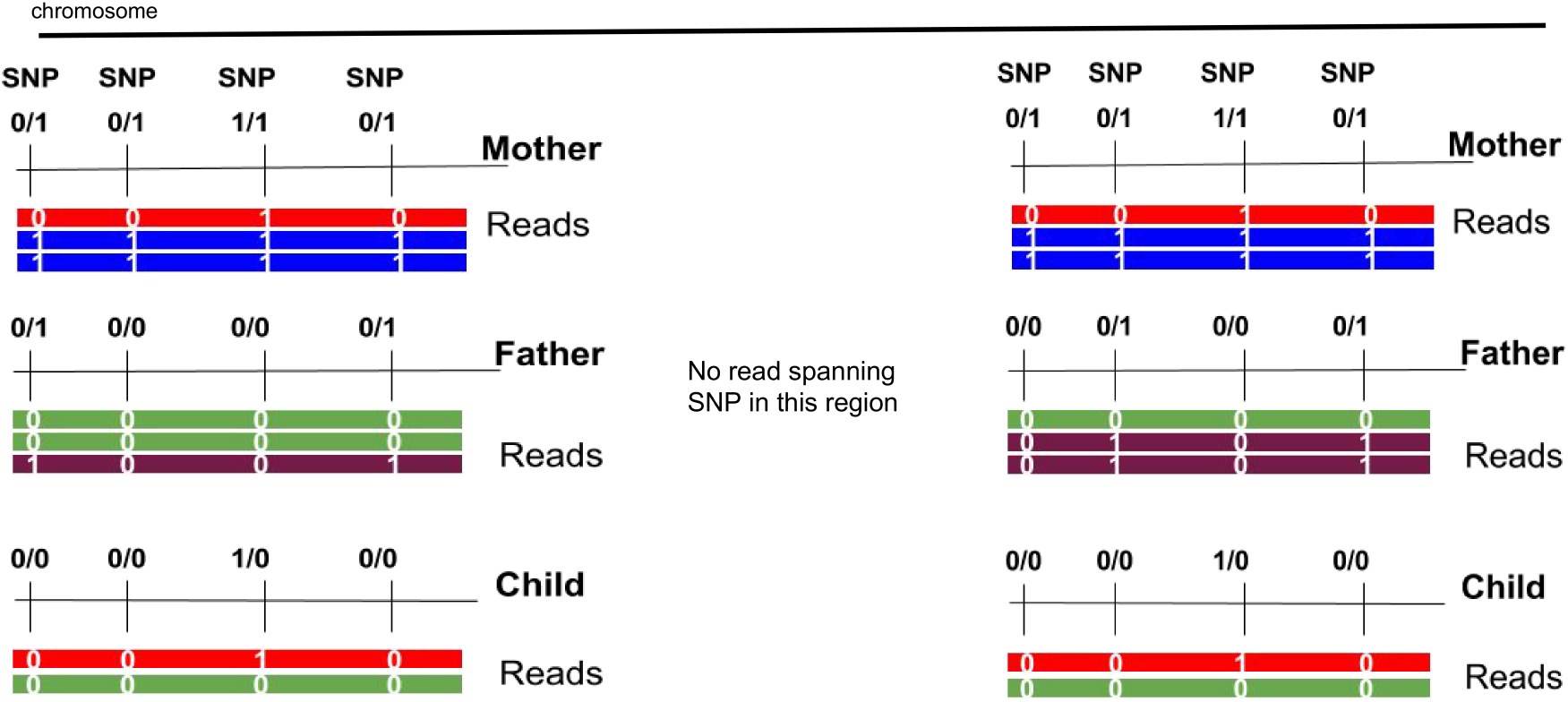
Two disjoint unconnected haplotype blocks for which phase information can be inferred from the genotypes.

### Runtimes

All experiments have been run on a server with two Intel Xeon E5-2670 CPUs (10 cores each) running at 2.5GHz. The implementation in WhatsHap is sequential, i.e. only using one CPU core. In all cases, the time spent reading the input files dominated the time spent in the phasing routine itself. Processing all three individuals of the 5× coverage real data set took 31.1 min, 31.2min, and 26.2min for wMEC-5, wMEC-15, and PedMEC-G-5, respectively. This time included all I/O and further processing. Of these times, 2.0 s, 4.1s, and 101.0 s were spent in the phasing routine, respectively. For input coverage 15x, total processing took 89.3 min, 93.9 min, and 65.4 min for wMEC-5, wMEC-15, and PedMEC-G-5, respectively. Of this, 2.5 s, 149.9 s, 321.5 s were spent in the phasing routines, respectively. We conclude that the phasing algorithm presented here is well suited for handling current data sets swiftly. In the future, we plan to further optimize the implementation of I/O subroutines and provide automatic chromosome-wise parallelization of data processing.

## 6 Discussion

We have presented a unifying framework for integrated read-based and genetic haplotyping. By generalizing the WhatsHap algorithm (Patterson *et al.*, 2015), we provide a fixed-parameter tractable method for solving the resulting NP-hard optimization problem, which we call *PedMEC*. When maximum coverage and number of individuals are bounded, the algorithm’s runtime is linear in the number of phased variants and independent of the read length, making it well suited for current and future long-read sequencing data. This is mirrored by the fact that the runtime is dwarfed by the time required for reading the input files in practice. PedMEC uses arbitrary costs for correcting errors in reads as well as for recombination events. By using phred-scaled probabilities as costs, minimizing the cost can be interpreted as finding a maximum likelihood phasing in a statistical model incorporating Mendelian inheritance, read error correction, and recombination.

Testing the implementation on simulated and real trio data, we could show that the method is notably more accurate than phasing individuals separately, especially at low coverages. Beyond enhanced accuracy, our method is also able to phase a greater fraction of heterozygous variants compared to single-individual phasing.

Being able to phase more variants is a key benefit of the integrative approach. Whereas read-based phasing can in principle only phase variants connected by a path through the covering reads, adding pedigree information enables even phasing of variants that are not covered at all since the algorithm can “fall back” to using genotype information. Figure 1 illustrates the increased connectivity while phasing a trio, resulting in more phased variants in practice.

Genetic haplotyping alone cannot phase variants that are heterozygous in all individuals, emphasizing the need for an integrative approach as introduced here. We demonstrate that such an approach indeed yields better result and recommend its use whenever both reads and pedigree information are available. Most remarkingly, the presented approach is able to deliver outstanding performance even for coverages as low as 2× per individual, on par with performance delivered by single-individual haplotyping at 15× coverage per individual.

### Future work

We plan to extend the implementation to support arbitrary pedigrees, although this extension would be harder to test since little suitable long-read data is available publicly. Phasing *de novo* variants arising in the child, which would be impossible with pure genetic haplotyping, is also straightforward to add.

Since runtime is exponential in the maximum physical coverage, pruning of datasets is required in practice. The read selection approach currently implemented in WhatsHap aims to retain reads both cover and connect many variants at the same time, in particular for heterogeneous combinations of datasets such as paired-end or mate-pair reads together with long reads. For pedigrees, each dataset is currently pruned individually, but results would likely improve if pedigree structure was taken into account in this step. Finally, since we show that it is possible and beneficial to integrate both read-based and genetic phasing, the next obvious question is whether it is possible to modify our unified theoretical framework to one that also includes statistical phasing.

## Acknowledgements

We thank Sarah O. Fischer for developing the read selection routine in WhatsHap.

## Funding

MM is supported by a grant from the Knut and Alice Wallenberg Foundation to the Wallenberg Advanced Bioinformatics Infrastructure.

## Footnotes

1 https://mathgen.stats.ox.ac.uk/impute/1000GP_Phase3.tgz.

2 http://www.shapeit.fr/files/genetic_map_b37.tar.gz

3 ftp://ftp-trace.ncbi.nlm.nih.gov/giab/ftp/data/AshkenazimTrio/(HG002_NA24385_son|HG003_NA24149_father|HG004_NA24143_mother)/PacBio_MtSinai_NIST/MtSinai_blasr_bam_GRCh37/

4 https://bitbucket.org/whatshap/whatshap

